# Human Rotavirus Diarrhea Is Associated with Altered Trafficking and Expression of Apical Membrane Transport Proteins

**DOI:** 10.1101/784264

**Authors:** Nicholas C. Zachos, Nicholas W. Baetz, Akshita Gupta, Anirudh Kapoor, Robert N. Cole, Alan S. Verkman, Jerrold R. Turner, Mary K. Estes, Mark Donowtiz

## Abstract

**Background:** Rotavirus (RV) is the 5^th^ leading cause of death in children <5 years old but the leading cause of diarrhea related deaths in this age group. The mechanism of RV diarrhea involves decreased activity of Na^+^-dependent solute transporters with increased luminal secretion of Cl^-^ in the absence of significant histologic damage. While our understanding of RV diarrhea has come from studies in animal models and cancer cell lines, the mechanism of the diarrhea and the transport proteins affected in human RV disease remains only partially understood. This understanding is likely to impact drug development therapy for RV diarrhea.

**Methods:** Formalin-fixed paraffin-embedded small intestinal specimens from patients diagnosed with RV diarrhea (confirmed by anti-RV antibodies) were analyzed by immunofluorescence for changes in apical/basolateral ion/nutrient transporters/channels as well as tight junctional and cytoskeletal proteins. Proximal small intestinal enteroids generated from biopsies obtained from healthy human subjects were grown as monolayers, differentiated to resemble villus epithelial cells, and infected with human RV.

**Results:** RV diarrhea was associated with reduced expression and intracellular localization of transport proteins normally found in the brush border membranes, including SGLT1, NHE3, NHE2, the Na^+^-dependent amino acid transporter SLC6A19, and CFTR. In contrast, basolateral proteins, including Na^+^/K^+^-ATPase, NKCC1, and β-catenin, the brush border marker ezrin, as well as the tight junction protein, ZO-1, were expressed and localized normally. RV-induced mislocalization of NHE3, SGLT1, SLC6A19 and CFTR was also seen when human small intestinal enteroids were infected with RV.

**Conclusions:** These data demonstrate a new pathophysiologic mechanism of acute diarrhea in which expression of multiple apical transport proteins are reduced. This acute diarrhea is likely to be caused by an effect on a common apical trafficking pathway, as exemplified by RV diarrhea, and its contribution to other enteric pathogen-induced diarrheal diseases should be determined.

## INTRODUCTION

In 2016, Rotavirus was the leading of diarrhea mortality in children <5 years old, causing ~128,000 deaths, as well as the leading cause of overall diarrhea mortality worldwide, causing ~228,000 deaths ^1^. The markedly reduced efficacy of oral RV vaccines in developing countries (39-49%), relative to the United States (90%), is one reason why overall RV-associated childhood death has fallen by only ~3% ^2, 3^. Oral rehydration solution (ORS) can be effective in RV diarrhea, but is used in <50% of cases. New approaches to RV diarrhea protection and therapy are, therefore, desperately needed.

RV preferentially infects differentiated enterocytes and enteroendocrine cells of the small intestine to induce a self-limiting, non-inflammatory diarrhea. The mechanisms identified as being involved in acute RV diarrhea include malabsorption, due to decreased enterocyte absorption, and increased fluid secretion induced by the RV enterotoxin, NSP4 ^4–6^. NSP4 is a virus-encoded ion channel that depletes Ca^2+^ from the ER of infected cells leading to stimulation of store operated Ca^2+^ entry to sustain elevated levels of intracellular Ca^2+^ that is critical for RV replication. RV transiently stimulates Cl^-^ secretion and since RV diarrhea occurs in models of cystic fibrosis, the Cl^-^ channel involved is unlikely to be CFTR ^7^. Moreover, in a mouse model of RV diarrhea, a Ca^2+^ activated Cl^-^ channel inhibitor prevented RV diarrhea ^8^. Activity of Na^+^-solute cotransporters, such as SGLT1, and the Na^+^/H^+^ exchanger, NHE3, are also reduced in RV ^9, 10^. Whether a common mechanism explains the changes in so many apical transport processes is not known. Despite studies using human colon cancer-derived cell lines and animal intestines (e.g. mice, rats, cats), there are no detailed studies using intestinal tissues from patients documented to have RV diarrhea. We and others have demonstrated that RV pathogenesis can be studied using human enteroids, which are self-propagating primary cultures of normal human small intestinal epithelium derived from intestinal stem cells obtained from healthy donors ^11^.

In this report, we analyzed pathology specimens of intestine from patients with documented RV diarrhea to describe a new mechanism for RV diarrhea that explains many of the previously reported effects on intestinal transport. We also demonstrate that a similar phenotype occurs in RV infected human enteroids, supporting future pathophysiologic studies in human enteroids and identifying a model to use for development of anti-RV diarrheal drugs.

## METHODS

### Identification of intestinal pathology specimens of cases of rotaviral diarrhea

A text for the term “rotaviral diarrhea” was applied to the University of Chicago Department of Pathology records, and formalin-fixed, paraffin-embedded (FFPE) intestinal biopsies were obtained. Similar numbers of disease-free biopsies from age (ages 2-22) and sex matched controls processed contemporaneously with the RV specimens were collected to serve as controls for processing and storage effects.

### Antibodies

Rabbit polyclonal RV antibodies against outer capsid protein, VP6 (1:50 dilution) and RV enterotoxin, NSP4 (1:100 dilution), have been previously described ^4^. Rabbit polyclonal antibody to: NHE3 (SLC9A3) was from Novus Biologicals (Cat #: NBP1-82574); NHE2 (SLC9A2)^14^; B°AT1 (SLC6A19) was from Sigma (Cat #: HPA043207); zonula occludins 1 (ZO-1) was from Invitrogen (Cat #: 33-9100); and β-catenin were from R&D Systems. Mouse monoclonal antibodies to: CFTR (clone M3A7; Cat #: 05-583) and SGLT-1 (Cat #: 07-1417) were from Millipore; NKCC1 (SLC12A2; Cat #: T4) and Na^+^/K^+^-ATPase (Cat #; a5) were from the University of Iowa Developmental Studies Hybridoma Bank. Rabbit monoclonal anti-phospho-ezrin (p-T567) was from Abcam (Cat #: EP2122Y).

### Human Rotavirus

Human RV strain, Ito, was replicated in MA104 cells, purified, and infected in human enteroids as previously described ^11^.

### Human enteroids

Human duodenal and jejunal enteroids were each established from four healthy donors (ages 2-22) obtained via endoscopic or surgical procedures utilizing the methods developed by the Clevers laboratory ^15^. De-identified biopsy tissue was obtained from healthy subjects who provided informed consent at Johns Hopkins University and all methods were carried out in accordance with approved guidelines and regulations. All experimental protocols were approved by the Johns Hopkins University Institutional Review Board (IRB#: NA_00038329 and IRB00044373). Briefly, enteroids generated from isolated intestinal crypts ^16^ were maintained as cysts embedded in Matrigel (Corning, USA) in non-differentiation media (NDM) containing Wnt3A, R-spondin-1, noggin and EGF, as we have described previously ^17^. Enteroid monolayers were generated as previously described in detail ^17, 18^. Monolayer differentiation was induced by incubation in Wnt3A-free^20^ and Rspo-1-free DFM for five days ^17^. Monolayer confluency and differentiation were monitored by measuring TER with an ohmmeter (EVOM^2^ World Precision Instruments, USA). The unit area resistances (ohm-cm^−2^) were calculated considering Transwell surface area (0.33 cm^2^) ^17, 18^.

### Immunofluorescence confocal microscopy

FFPE blocks were sectioned to 5μm and rehydrated through an ethanol gradient prior to microwave antigen retrieval in 10mM sodium citrate (pH 6.0). For enteroids, monolayers were fixed in 2% paraformaldehyde in PBS for 30 minutes. Sections and monolayers were blocked and permeabilized in PBS containing 2% BSA, 15% FBS, and 0.1% saponin for 1 hour and then washed three times in PBS. Primary antibodies (1:100 dilution) in PBS were incubated overnight at 4°C followed by three PBS washes. Cells were then incubated with anti-mouse or anti-rabbit AlexaFluor (488nm or 568nm) conjugated secondary antibodies (diluted 1:100) and Hoescht 33342 (nuclear counterstain) in PBS for 1 hour at room temperature. Tissue sections and enteroid monolayers were washed in PBS, mounted, and coverslipped. Confocal images were obtained on Zeiss 510 META confocal microscope. The Atlas of Intestinal Transport (https://www.jrturnerlab.com/atlasofintestinaltransport) was used to assess localization of specific transport proteins in small intestinal segments from human subjects.

### LC-MS/MS analysis

Peptides in 28 fractions after basic reverse phase (bRP) fractionation, dried, 8.57ug/fraction, were re-constituted in 12uL 2%ACN/0.1%FA and 6uL, ~4.25ug (50%) analyzed by liquid chromatography/tandem mass spectrometry (LC-MS/MS) in FTFT using QExactive Plus (Thermo Fisher Scientific) interfaced with nano-Acquity LC system from Waters. Peptides were loaded on a 75 um x 2.5 cm C18 (YMC*GEL ODS-A 12nm S-10 μm) trap at 600nl/min 0.1% FA (solvent A) and fractionated at 300 nL/min on a 75 um x 100 mm ProntoSil C18H reverse-phase column (5 μm, 120Å, http://www.bischoff-chrom.com/hplc-prontosil-c18-h-c18-phasen.html) using a 2-10% solvent B (90% acetonitrile in 0.1% formic acid) gradient over first 2min, then up to 25% B by 55min, 45% B by 67 min and 100% B by 75min. Eluting peptides were sprayed into an LTQ Orbitrap Velos mass spectrometer through 1 μm emitter tip (New Objective, www.newobjective.com) at 2.4 kV. Survey scans (full ms) were acquired on Orbi-trap within 350-1800Da m/z using Data-dependent Top 12 method with dynamic exclusion of 30 s. Precursor ions were individually isolated with 1.2Da, fragmented (MS/MS) using HCD activation collision energy 30. Precursor and the fragment ions were analyzed at resolution (at 200Da) 70,000 and 35000, respectively.

### Data Analysis

MS/MS spectra were analyzed via Proteome Discoverer software (v1.4 Thermofisher Scientific) using 3Nodes (extracted, processed by MS2Processor and PD1.3), with Mascot (v 2.5,1 Matrix Science, London, UK) using the RefSeq2015 Complete Database with concatenated decoy database specifying human species, Trypsin/P as enzyme, missed cleavage 2, fixed modifications TMT10-plex on N-termini and carbamidomethylation on cysteine, and variable modifications include TMT 10plex labeling of lysines, oxidation of methionine and deamidation on Precursor tolerance 12ppm, fragment tolerance 0.03Da, extract resolution 55K at 400Da. NQ. Peptide identifications from Mascot searches were processed within the Proteome Discoverer to identify peptides with a confidence threshold 0.01% False Discovery Rate, based on a concatenated decoy database search and to calculate the protein and peptide ratios.

### Grant Support

The current study was funded by NIH grants: R24-DK099803 (MD), R01-AI080656 (ME), R01-DK026523 (MD), P30-DK089502 (MD, NZ), R03-DK091482 (NZ), P30-DK72517 (AV), and R01-DK061931 (JRT).

Funding source NIH grants had no influence in study design, analysis, data interpretation or writing of the manuscript.

## RESULTS

### Validation of RV infection in pathology specimens

Cases previously diagnosed as rotaviral diarrhea and controls were analyzed by immunofluorescence confocal microscopy. Of 17 cases initially categorized as RV diarrhea, 12 cases were confirmed positive for rotaviral infection on the basis of immunofluorescent staininng for both VP6 and NSP4. Neither protein was detected in any of the the healthy controls (**Fig 1**). These 12 cases were studied further.

**FIGURE 1:**
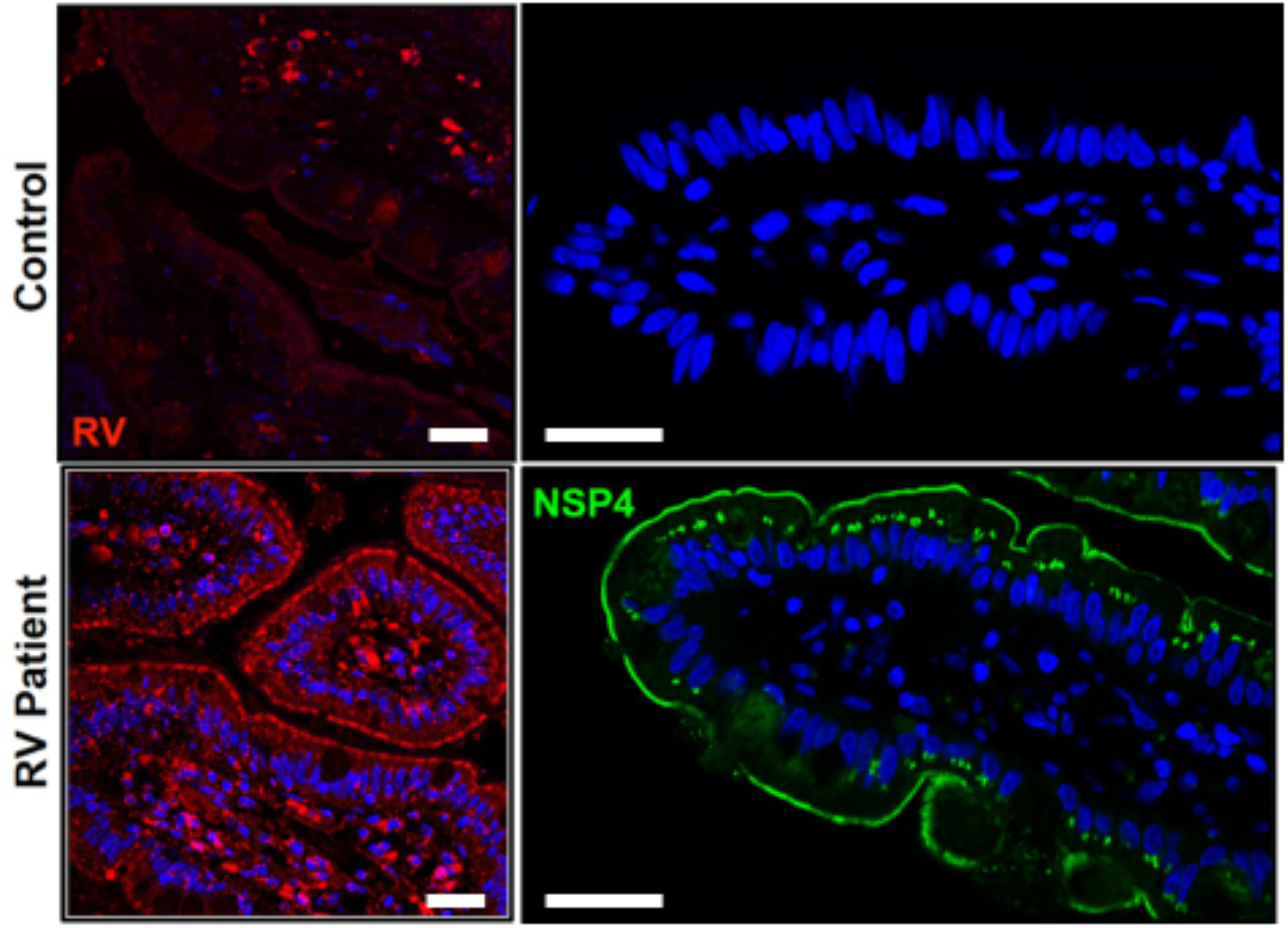
RV antigen and NSP4 were detected in all human intestinal rotavirus pathology specimens further studied. Immunofluorescent detection of RV (red) and NSP4 (green) in pathology specimens of the small intestine from patients documented with RV diarrhea (bottom) and aged-matched healthy subjects (control; top). RV and NSP4 expressed at the BB and intracellularly in villus epithelial cells. Scale bar = 20μm.

### RV diarrhea is associated with altered localization and/or expression of brush border ion/nutrient transporters and channels

Immunofluorescence confocal microscopy revealed major differences in brush border/apical transport proteins exhibiting mislocalization away from the brush border as well as reduced expression in RV specimens compared to healthy controls. This was true of NHE2, NHE3, CFTR, SGLT1, and B°AT1 (SLC6A19) (**Fig 2A, B and Table 1**). SGLT1 localization was subapical while CFTR, NHE2 and SLC6A19 were redistributed towards the basolateral pole of epithelial cells. The overall expression of NHE3 in enterocytes was nearly lost. In some cases, expression of NHE2, SGLT1 or CFTR could also not be detected (**Fig 2A**). In order to confirm that these changes were not due to general structural changes in the brush border, which appeared normal by histologic analysis, we examined expression and localization of phosphorylated (T567) ezrin (i.e. p-ezrin), a brush border structural protein, as well as tight junction (TJ) markers ZO1 and occludin. The location and magnitude of p-ezrin and ZO-1 expression were not affected rotavirus infection, relative to healthy controls (**Fig 3, Table 1**). To consider whether there was a general abnormality of transport protein expression or localization, the basolateral transport proteins NKCC1 and Na^+^/K^+^-ATPase as well as β-catenin (data not shown) were studied. No changes in these proteins were identified (**Figs 3, 4, Table 1**). Thus, RV specifically impairs the apical expression of multiple small intestinal ion and nutrient transporters and channels without affecting basolateral transporters or brush border and TJ structural proteins.

**FIGURE 2:**
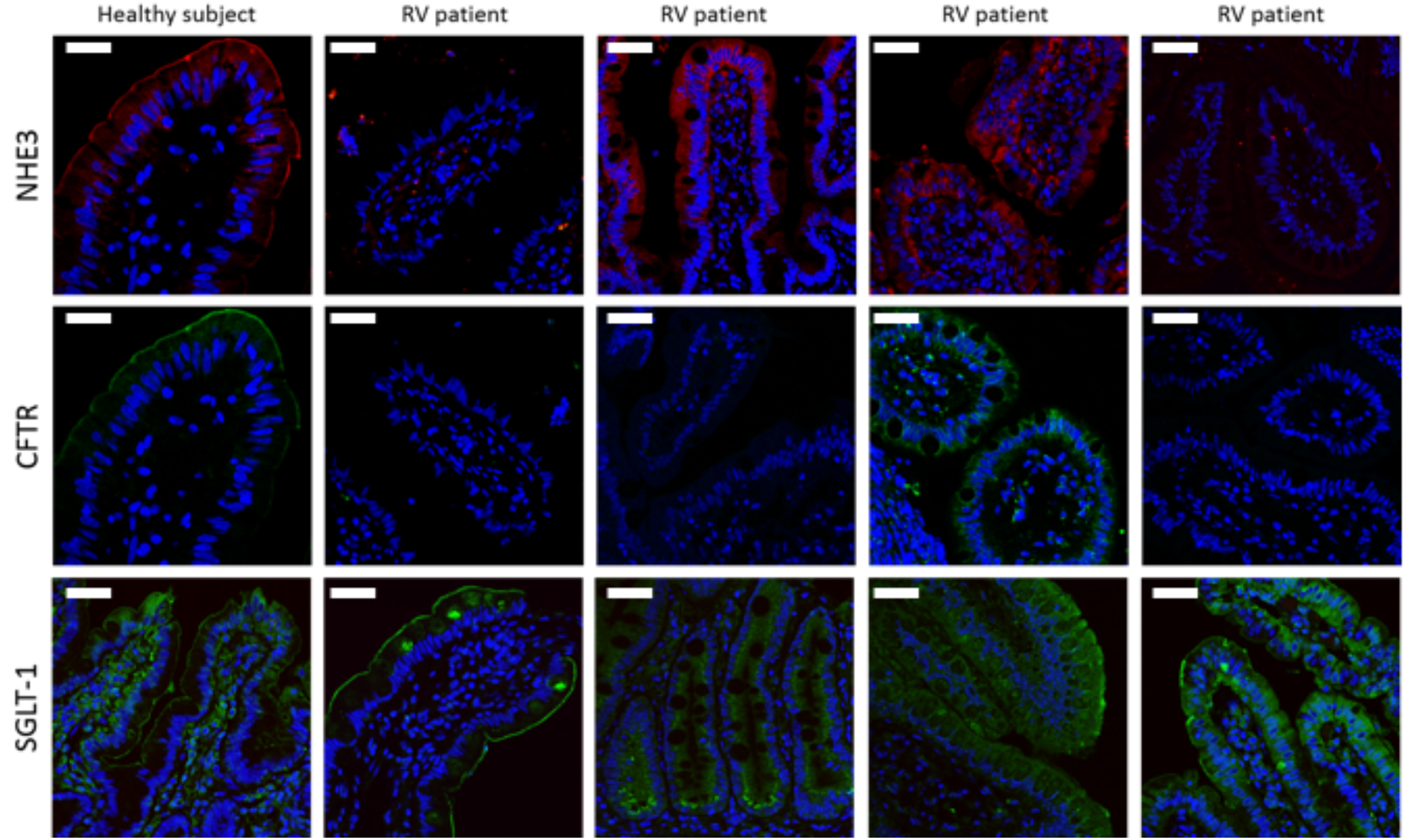
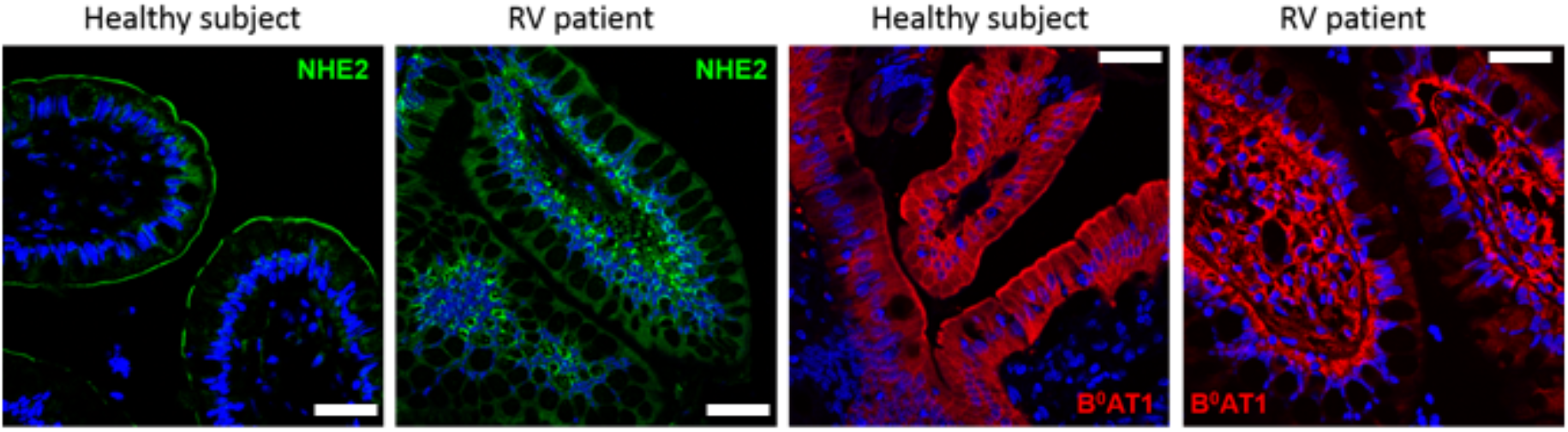
BB localization and expression of apical ion/nutrient transporters is altered in the small intestine of RV patients compared to age-matched healthy subjects. FFPE patient small intestinal specimens were immunostained for NHE2 (green), NHE3 (red), CFTR (green), SGLT1 (green), and B^0^AT1 (red) and images obtained by confocal microscopy. Blue = nuclei. Scale bar = 20μm.

**TABLE 1:** Summary of RV effects on intestinal transporters in RV infected patient specimens.

| Antibody | Normal presentation | Expression/Localization |
| --- | --- | --- |
| CFTR | 0/12 | Downregulated/BLM |
| NHE2 | 2/12 | Downregulated/Cytosolic |
| NHE3 | 4/12 | Downregulated/Cytosolic |
| SGLT1 | 3/12 | Downregulated/Cytosolic |
| B<sup>0</sup>AT1 | 3/11 | No BB; Basal concentration |
| p-Ezrin | 12/12 |  |
| Na<sup>+</sup>/K<sup>+</sup> ATPase | 11/11 |  |
| NKCC1 | 11/11 |  |

**FIGURE 3:**
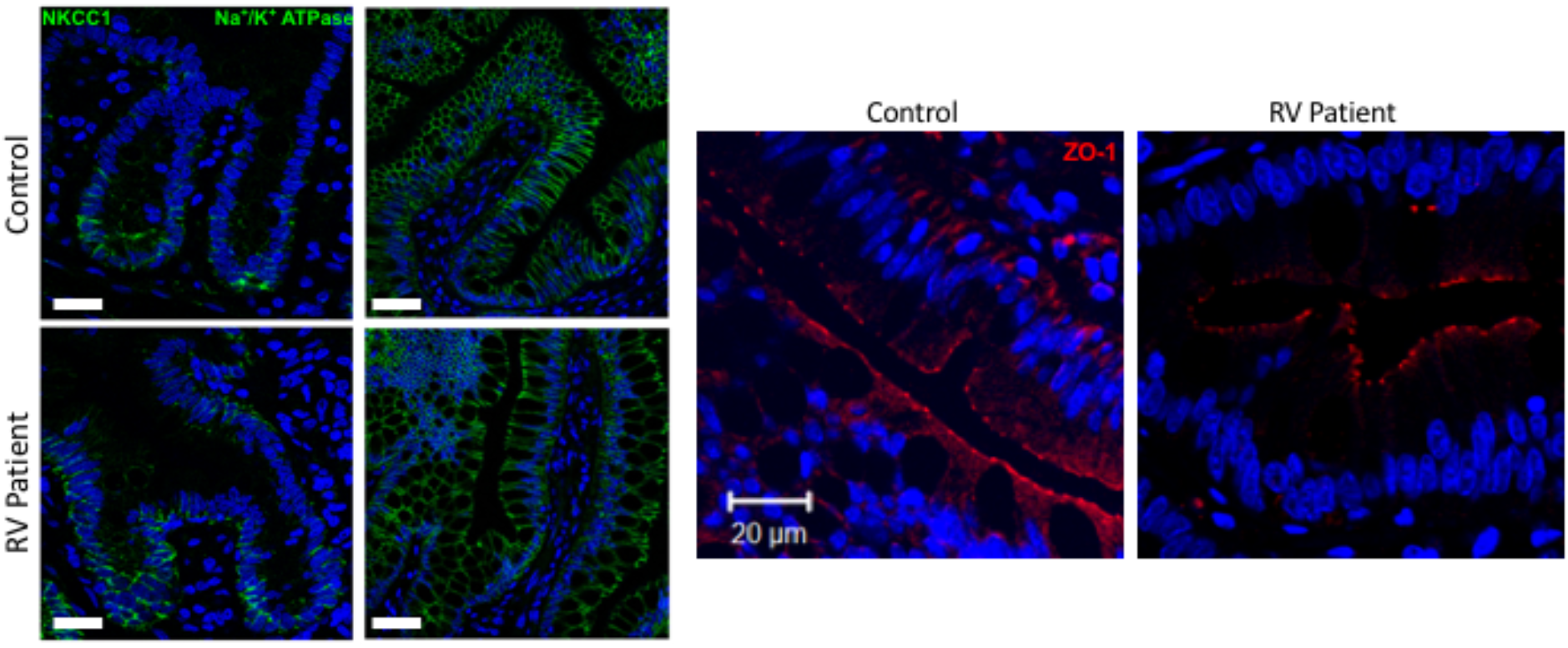
BB localization and expression of phosphorylated ezrin (p-ezrin) is similar in the small intestine of RV patients compared to age-matched healthy subjects. Small intestinal specimens from healthy subjects and RV infected patients were immunostained for the apical cytoskeletal marker, phosphor-ezrin (p-ezrin; T567) and images captured using confocal microscope. Blue = nuclei. Scale bar = 20μm.

**FIGURE 4:**
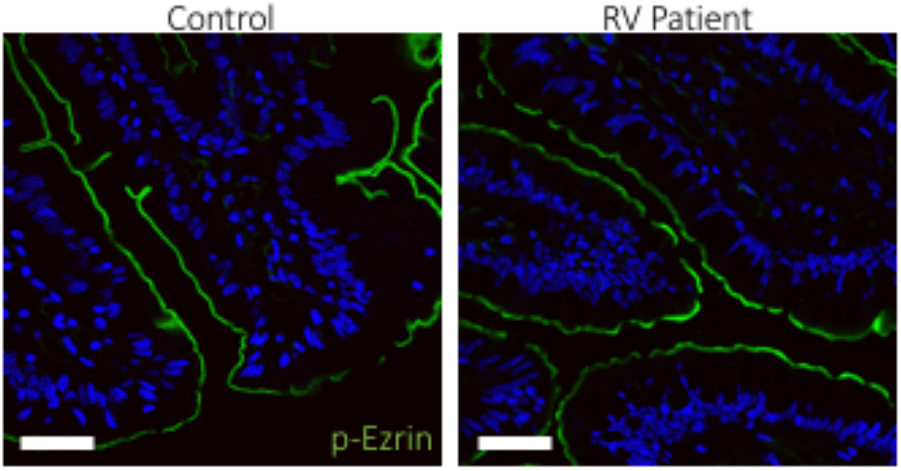
Expression and basolateral localization of NKCC1 and Na^+^/K+ ATPase (each in green on left) and TJ location of ZO1 (red on right) are not affected in RV cases. All images: blue = nuclei. Scale bar = 20μm.

### Mislocalization or downregulation of apical transporters present in other diarrhea diseases

In addition to profiling the expression and localization of intestinal transporters/channels and tight junctional proteins in healthy subjects, The Atlas of Intestinal Transport is also comprised of FFPE tissues sections obtained from healthy subjects and patients with various types of intestinal disorders that have diarrhea as a common symptom. Using this Atlas, we analyzed whether RV-mediated changes in the expression of apical transporters also occur in other types of diarrheal disease. Examination of tissue sections from some patients with villous atrophy or common variable immunodeficiency revealed decreased expression of NHE3 and/or partial mislocalization of CFTR and SGLT-1. Similar to RV patient specimens and healthy controls, the expression and localization of ZO-1 appeared normal as did the basolateral marker, E-cadherin. These findings support the hypothesis that some diarrheal diseases, in addition to acute RV diarrhea, are associated with altered expression/localization of apical transport proteins.

### Infection of human enteroids with human RV strain recapitulates the changes in brush border/apical domain localization of multiple transport proteins

Human RV strains can infect adult intestinal crypt-derived enteroids from each segment of human small intestine as well as replicate over at least 96h as detected by quantitation of increasing viral RNA by qRT-PCR, virus-specific protein expression by Western blot analysis, and production of infectious virus by fluorescent focus assays ^11^. Laboratory strains of human RV as well as human clinical isolates can replicate in enteroids, which subsequently produce infectious rotavirus ^11^. Whether RV infection of differentiated (5 days after Wnt3A, R-spondin-1 removal) human small intestinal enteroids altered apical transporter expression or localization was determined by exposure to human RV with mock infected enteroids as negative controls, as described previously ^11^. The conditions studied were associated with RV infection (~40% infected cells similar to previously shown ^11^) as indicated by presence of virus visualized by transmission electron microscopy within 1 hour post infection (**Fig 5**). Monolayers were fixed at multiple times after RV infection (1,6,24h) and immunostained with the same antibodies against intestinal transporters used to analyze RV patient specimens. **Figures 6 and 7** demonstrate by immunofluorescence confocal microscopy that RV infection reduced the surface expression of CFTR, NHE3, and SGLT-1 similarly to that observed in patient samples. At 6 hours post infection, CFTR was miss-localized to the basolateral membrane compared to mock infected controls in which CFTR was exclusively localized to the apical domain (**Fig 6**). By 24 hours post infection, SGLT-1 and NHE3, which are normally expressed in the apical membrane of mock infected enteroid monolayers, exhibited increased expression in the cytosol below the apical membrane, with NHE3 expression markedly reduced (**Fig 7**). These results demonstrate that effects of human RV infection on the surface expression of apical ion/nutrient transporters in human small intestinal enteroids mimics changes in clinical cases of RV infection.

**FIGURE 5:**
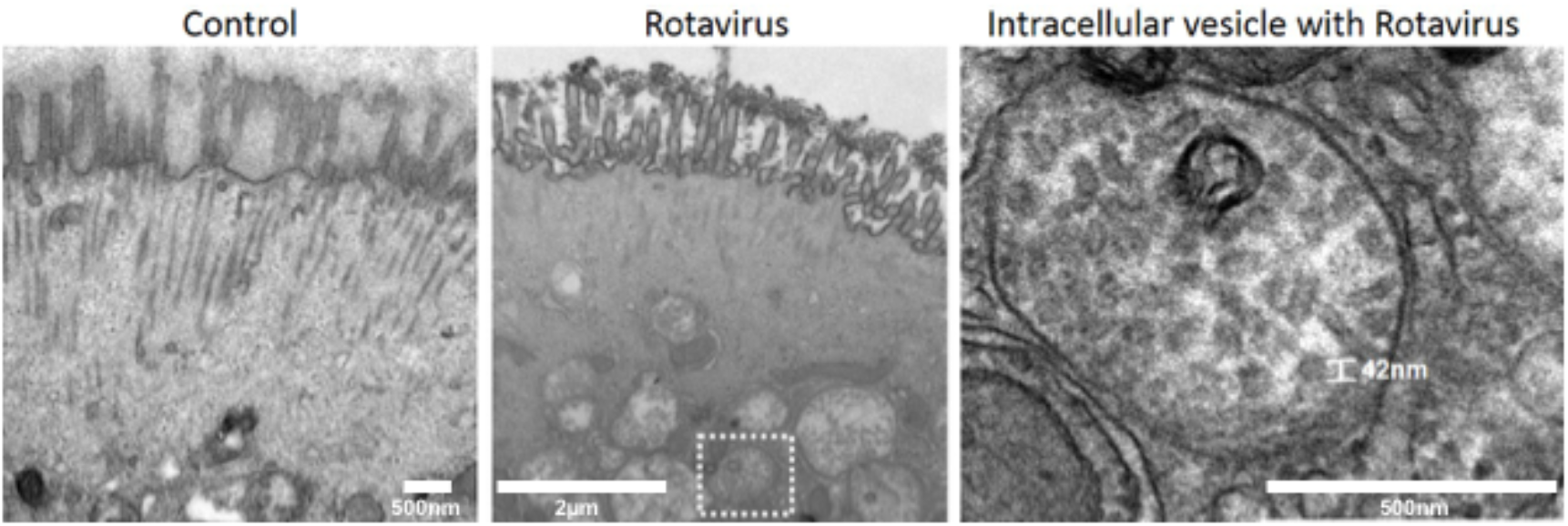
RV particles detected in large, non-electron dense, intracellular vesicles in human enteroids. RV particles (~40-50nm) were observed at 1 hour post infection in intracellular vesicles ranging from 200nm-1μm in diameter. Microvilli exhibit partial blunting following RV infection. *Right:* Higher magnification of internalized RV particles from region denoted by white dash box.

**FIGURE 6:**
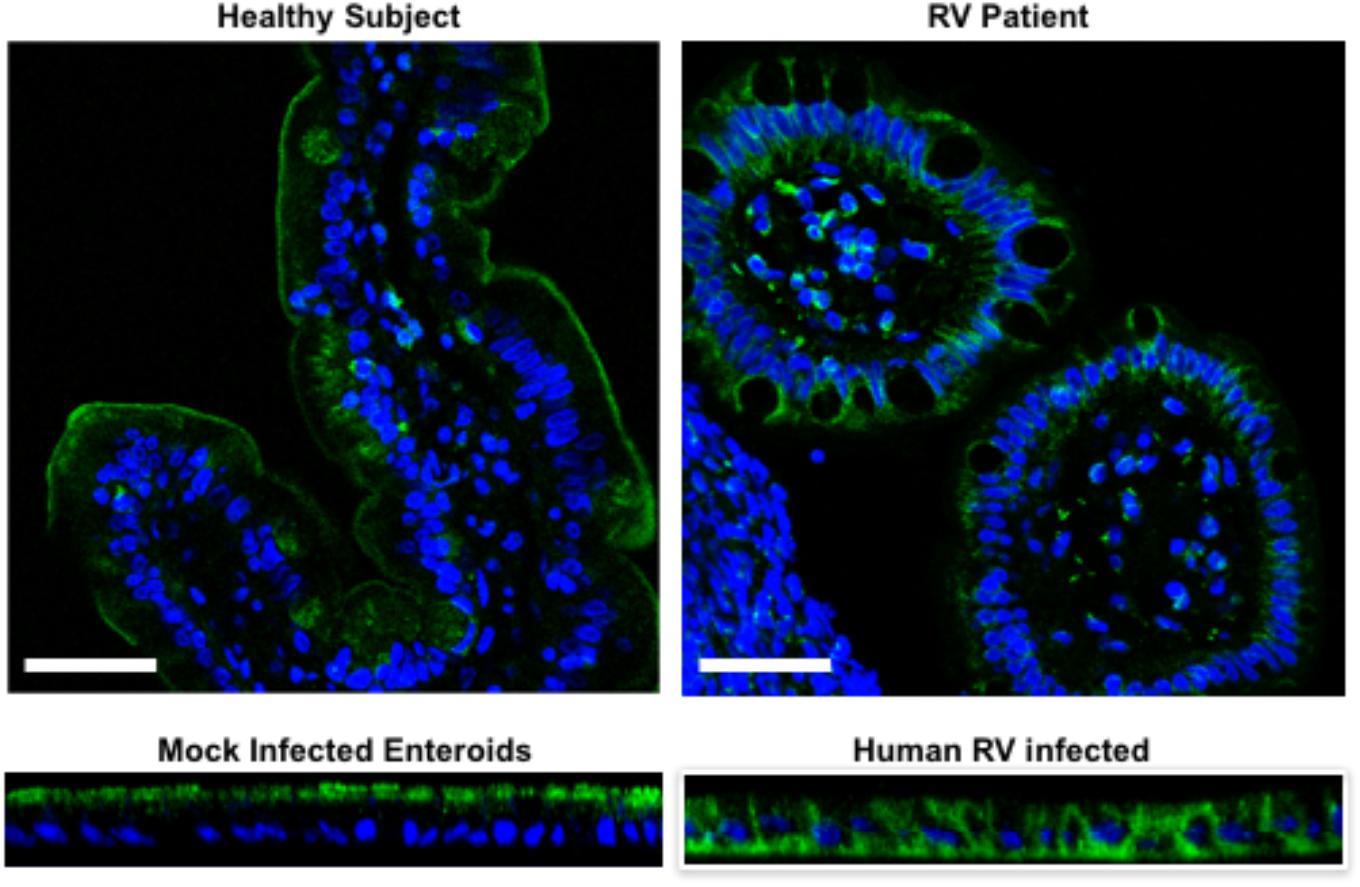
Enteroids mimic CFTR miss-localization in RV patient specimens. Following RV infection in human enteroids, monolayers were fixed in 2% PFA and immunostained for CFTR. Confocal microscopy was used to capture optical sections and the resulting orthogonal planes are presented with apical membrane at the top of each image. Blue = nuclei.

**FIGURE 7:**
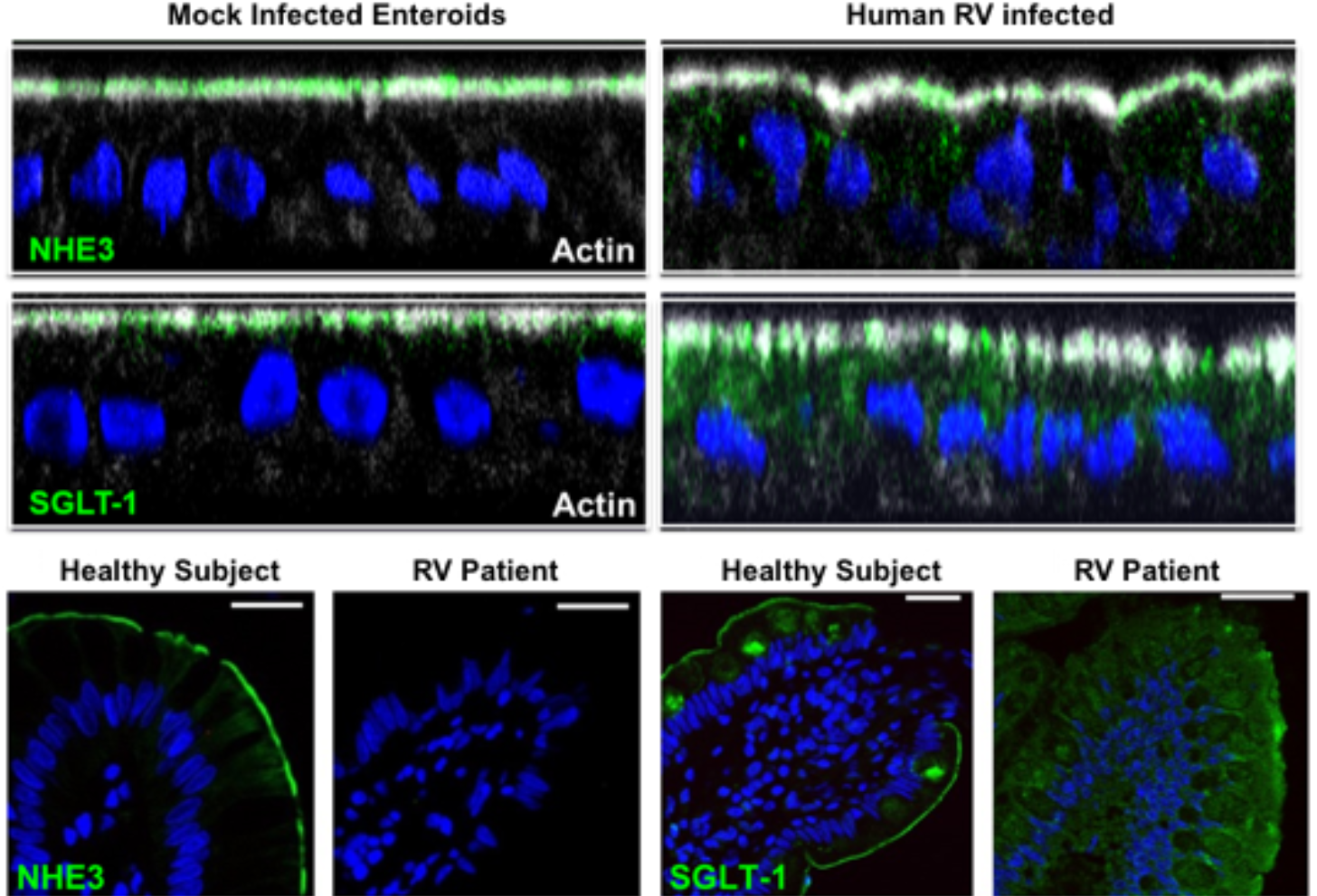
Enteroids mimic NHE3 and SGLT-1 miss-localization in RV patient specimens. Human enteroids infected with RV were fixed after 24 hours and immunostained for NHE3 and SGLT1 (green). Optical sections were obtained by confocal microscopy and orthogonal images (below) represent a cross sectional view of enteroid monolayers with the apical membrane at the top of each image. Blue = nuclei.

### Altered expression and localization of apical ion/nutrient transporters is associated with decreased expression of proteins associated with apical endosome sorting and/or recycling

In order to characterize changes in protein trafficking due to RV infection, we performed quantitative proteomics on human small intestinal enteroid monolayers mock or RV infected for 1-48 hours. As summarized in **Table 2**, RV significantly downregulated expression of multiple apical trafficking proteins including the apical cargo trafficking motor protein, myosin Vb, and Ras associated binding (Rab) proteins, and associated binding partners that regulate cargo trafficking via the apical endosome.

**TABLE 2:**
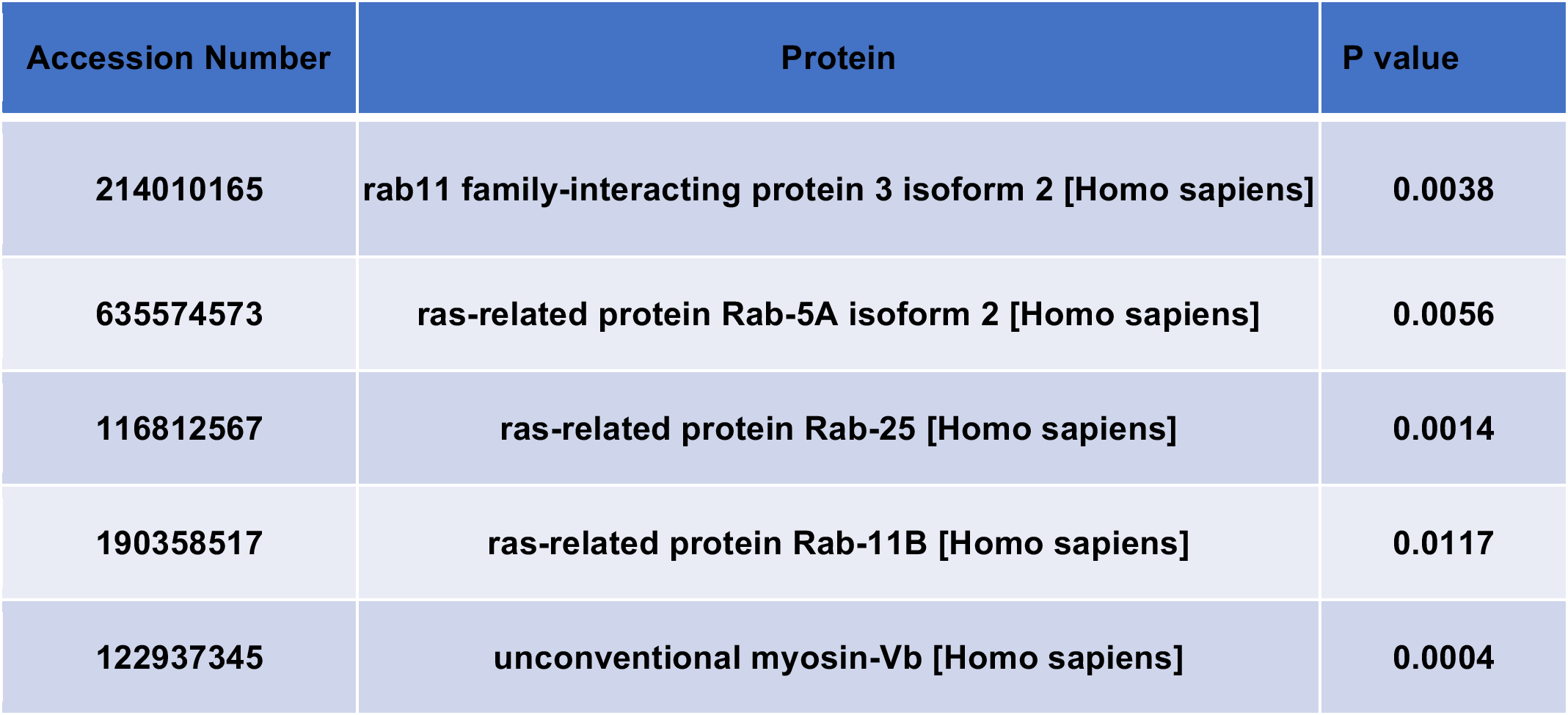
Proteomics analysis of human small intestinal enteroids infected with RV.

| Accession Number | Protein | P value |
| --- | --- | --- |
| 214010165 | rab11 family-interacting protein 3 isoform 2 [Homo sapiens] | 0.0038 |
| 635574573 | ras-related protein Rab-5A isoform 2 [Homo sapiens] | 0.0056 |
| 116812567 | ras-related protein Rab-25 [Homo sapiens] | 0.0014 |
| 190358517 | ras-related protein Rab-11B [Homo sapiens] | 0.0117 |
| 122937345 | unconventional myosin-Vb [Homo sapiens] | 0.0004 |

## DISCUSSION

The findings of this study define the impact of RV infection on the expression and localization of the major intestinal epithelial ion/nutrient transporters/channels in human intestinal tissue. We observed that multiple apical transport proteins, including Na^+^-dependent ion/solute transporters and the major intestinal chloride secretory channel, CFTR, are either mislocalized and, in some cases, downregulated in the small intestine of patients with RV diarrhea when compared to specimens from healthy subjects. These changes in expression/localization appeared to be specific for apical transporters/channels since basolateral transporters, as well as tight junctional proteins, were normally distributed in RV patient specimens. Moreover, the changes observed in clinical specimens can be reproduced in a primary human *in vitro* epithelial culture system (i.e. enteroids). Due to the specific effects of RV infection on apical transporters, our data suggest that a common apical trafficking pathway shared by multiple transporters/channels may be the mechanism responsible for RV diarrhea.

Previous studies of RV diarrhea have separately addressed RV effects on individual transporters. Together, these data have suggested the RV diarrhea occurs due to increased Cl^-^ secretion and defective Na^+^ absorption. The results of our study unify previous findings by examining how RV alters the expression/localization of multiple transporters in human enteroids and comparing these phenotypes to clinical specimens from RV infected patients. Here we show that RV diarrhea appears to be due to mild changes in the brush border, seen by EM but not by light microscopy and not associated with changes in expression of the brush border protein p-ezrin or the tight junction protein ZO-1, plus abnormal apical membrane trafficking of transport proteins. The abnormal trafficking of the proteins identified in this study, would account for the documented reduced Na^+^ and glucose absorption (reduced apical SGLT1, NHE3), occurrence of diarrhea in CF models (CFTR not apically localized), and ability of the calcium activated Cl^-^ channel inhibitor ^8^ to totally prevent RV diarrhea in a neonatal mouse model. However, the calcium activated Cl^-^ channel involved in small intestinal secretion has not been identified molecularly. Consequently, we cannot determine if that protein is also mislocalized or down-regulated; although, the results suggest it is trafficked differently than the affected proteins. The fact that ORS is marginally effective in rehydrating patients with RV diarrhea is at least partially explained by the fact that RV infection reduces the overall amount of transporters that could respond to ORS. However, the fact that ORS saves lives in RV diarrhea speaks for the fact that the transport processes involved in ORS related rehydration (SGLT1, NHE3, Na^+^/K^+^-ATPase) are still at least partially functional, which is consistent with the previous documentation of patchy histologic changes in small intestine of RV diarrhea.

The current report potentially identifies a previously unrecognized mechanism of diarrhea that might be relevant for additional difficult to treat diarrheas; however, the precise mechanism involved abnormal apical trafficking remains to be determined. To support the hypothesis that apical trafficking is the general mechanism responsible, we identified a panel of apical endosome associated proteins that are significantly downregulated following RV infection in human enteroids (Table 2) and have known regulatory effects on apical transporters. Future studies will be necessary to demonstrate that a loss of these proteins can phenocopy the expression and/or localization of changes observed after RV infection. In a parallel report [Maeda and Zachos et al], we described that RV infection of intestinal epithelial cells was associated with reduced expression of PARD6B, a protein known to be involved in apical recycling ^19^. This RV effect was associated with increased proteasomal degradation of PARD6B as well as atypical PKC, which is part of the PARD6B polarity complex. Moreover, gene suppression of PARD6B expression in MDCK, Caco-2 cells, and human enteroids disrupted apical endosomal trafficking and altered expression of apical membrane transport proteins, similar to what we observed in RV patient specimens. It is not known whether these RV effects account for the reduced expression of apical transport proteins demonstrated in the human pathology specimens. Taken together, both studies suggest that RV mediates the altered expression and localization of apical ion/nutrient transporters by affecting components of the apical endosome. These results may be potentially important for directing development of medical management of acute rotaviral diarrhea, since developing drugs to stimulate NHE3 or inhibit CFTR, which are the commonly suggested targets of anti-diarrheal drug development, are not likely to be successful.

Importantly, the reproduction in human enteroids of the RV effects on apical trafficking and transport protein expression in the human disease provides a model for additional mechanistic studies and potential drug development to treat RV diarrhea. We recently established the methodology to grow human enteroids as confluent 2D monolayers to generate a controllable and tractable model to study luminal host-pathogen interactions and standardized this model to allow separate study of a “villus”-like epithelium (differentiated enteroids) and “cryp”-like epithelium (undifferentiated enteroids) ^17, 18^. Furthermore, we showed that RV preferentially targets differentiated enteroids, which is consistent with the previous recognition that RV preferentially infects villus tip enterocytes ^20^. We further established that human enteroids demonstrate intracellular changes typical of RV replication in cultured cells, including visualization of nascent virus particles within intracellular vesicles by electron microscopy (**Figure 4**), some microvillar damage, the induction of lipid droplets, and formation of replication compartments ^11^.

While RV is the first diarrheal disease to act by causing a defect in the general apical trafficking mechanism, it will be important to determine how wide-spread is the contribution of this mechanism to other diarrheal diseases and to diseases of other epithelia (lung, kidney, brain, etc). Understanding what transport processes are abnormal in diarrheal diseases and the mechanisms by which the changes occur, provides strategies for drug development in an area that currently lacks effective treatments.

## REFERENCES

1. Tate JE, Burton AH, Boschi-Pinto C, Parashar UD; World Health Organization–Coordinated Global Rotavirus Surveillance Network. Global, Regional, and National Estimates of Rotavirus Mortality in Children <5 Years of Age, 2000-2013. Clin Infect Dis. 2016 May 1;62 Suppl 2:S96–S105.

2. Armah GE, Sow SO, Breiman RF, Dallas MJ, Tapia MD, Feikin DR, et al. Efficacy of pentavalent rotavirus vaccine against severe rotavirus gastroenteritis in infants in developing countries in sub-Saharan Africa: a randomised, double-blind, placebo-controlled trial. Lancet. 2010;376(9741):606–14.

3. Zaman K, Dang DA, Victor JC, Shin S, Yunus M, Dallas MJ, et al. Efficacy of pentavalent rotavirus vaccine against severe rotavirus gastroenteritis in infants in developing countries in Asia: a randomised, double-blind, placebo-controlled trial. Lancet. 2010;376(9741):615–23.

4. Ball JM, Tian P, Zeng CQ, Morris AP, Estes MK. Age-dependent diarrhea induced by a rotaviral nonstructural glycoprotein. Science. 1886; 272: 101–104.

5. Halaihel N, Lievin V, Alvarado F, Vasseur M. Rotavirus infection impairs intestinal brush-border membrane Na(+)-solute cotransport activities in young rabbits. Am J Physiol Gastrointest Liver Physiol. 2000;279(3):G587–96.

6. Halaihel N, Lievin V, Ball JM, Estes MK, Alvarado F, Vasseur M. Direct inhibitory effect of rotavirus NSP4(114-135) peptide on the Na(+)-D-glucose symporter of rabbit intestinal brush border membrane. J Virol. 2000;74(20):9464–70. PMCID: 112375.

7. Morris AP, Scott JK, Ball JM, Zeng CQ, O’Neal WK, Estes MK. NSP4 elicits age-dependent diarrhea and Ca(2+)mediated I(-) influx into intestinal crypts of CF mice. Am J Physiol. 1999;277(2 Pt 1):G431–44.

8. Ko EA, Jin BJ, Namkung W, Ma T, Thiagarajah JR, Verkman AS. Chloride channel inhibition by a red wine extract and a synthetic small molecule prevents rotaviral secretory diarrhoea in neonatal mice. Gut. 2014;63(7):1120–9.

9. Foulke-Abel J, In J, Kovbasnjuk O, Zachos NC, Ettayebi K, Blutt SE, et al. Human enteroids as an ex-vivo model of host-pathogen interactions in the gastrointestinal tract. Exp Biol Med (Maywood). 2014;239(9):1124–34.

10. Zou WY, Blutt SE, Zeng XL, Chen MS, Lo YH, Castillo-Azofeifa D, Klein OD, Shroyer NF, Donowitz M, Estes MK. Epithelial WNT Ligands Are Essential Drivers of Intestinal Stem Cell Activation. Cell Rep. 2018 Jan 23;22(4):1003–1015.

11. Saxena K, Blutt SE, Ettayebi K, Zeng X-L, Broughman JR, Crawford SE, Karandikar U, Sastri NP, Conner ME, Opekun A, Graham DY, Foulke-Abel J, In J, Kovbasnjuk O, Zachos NC, Donowitz M, Estes MK. Human intestinal enteroids: a new model to study human rotavirus infection, host restriction and pathophysiology. Journal of Virology. 2015; 90(1): 43–56.

12. Blutt SE, Matson DO, Crawford SE, Staat MA, Azimi P, Bennett BL, Piedra PA, Conncer ME. Rotavirus antigenemia in children is associated with viremia. PLoS Med. 2007; 4: e121.

13. Sastri NP, Viskoyska M, Hyser JM, Tanner MR, Horton LB, Sankaran B, Prasad BV, Estes MK. Structural plasticity of the coiled-coil domain of rotavirus NSP4. J Virol. 2014; 88:13602–13612.

14. Hoogerwerf WA, Tsao SC, Devuyst O, Levine SA, Yun CH, Yip JW, Cohen ME, Wilson PD, Lazenby AJ, Tse CM, Donowitz M. NHE2_and NHE3 are human and rabbit intestinal brush-border proteins. Am J Physiol. 1996 Jan;270(1 Pt 1):G29–41.

15. Sato T, Vries RG, Snippert HJ, van de Wetering M, Barker N, Stange DE, et al. Single Lgr5 stem cells build crypt-villus structures in vitro without a mesenchymal niche. Nature. 2009;459(7244):262–5.

16. Sato T, Stange DE, Ferrante M, Vries RG, Van Es JH, Van den Brink S, Van Houdt WJ, Pronk A, Van Gorp J, Siersema PD, Clevers H. Long-term expansion of epithelial organoids from human colon, adenoma, adenocarcinoma, and Barrett’s epithelium. Gastroenterology. 2011 Nov;141(5):1762–72.

17. Noel GN, Baetz NW, Staab JF, Donowitz M, Kovbasnjuk O, Pasetti M, Zachos NC. A primary human macrophage-enteroid co-culture model to investigate mucosal gut physiology and host-pathogen interactions. Nature Scientific Reports. 2017; 7: 45270.

18. In J, Foulke-Abel J, Zachos NC, Hansen AM, Kaper JB, Bernstein HD, et al. Enterohemorrhagic reduce mucus and intermicrovillar bridges in human stem cell-derived colonoids. Cell Mol Gastroenterol Hepatol. 2016;2(1):48–62 e3. PMCID: 4740923.

19. Nelms B, Dalomba NF, Lencer W. A targeted RNAi screen identifies factors affecting diverse stages of receptor-mediated transcytosis. J Cell Biol. 2017 Feb;216(2):511–525.

20. Greenberg HB, Estes MK. Rotaviruses: from pathogenesis to vaccination. Gastroenterology. 2009;136(6):1939–51.

